# Impaired hand dexterity function in a non-human primate model with chronic Parkinson’s disease

**DOI:** 10.1101/2020.05.01.072058

**Authors:** Jincheol Seo, Jinyoung Won, Keonwoo Kim, Junghyung Park, Hyeon-Gu Yeo, Yu Gyeong Kim, Seung Ho Baek, Hoonwon Lee, Chang-Yeop Jeon, Won Seok Choi, Sangil Lee, Ki Jin Kim, Sung-Hyun Park, Yeonghoon Son, Kang Jin Jeong, Kyung Seob Lim, Philyong Kang, Hwal-Yong Lee, Hee-Chang Son, Jae-Won Huh, Young-Hyun Kim, Yeung Bae Jin, Dong-Seok Lee, Sang-Rae Lee, Ji-Woong Choi, Youngjeon Lee

## Abstract

Symptoms of Parkinson’s disease (PD) caused by loss of dopaminergic neurons are accompanied by movement disorders, including tremors, rigidity, bradykinesia, and akinesia. Non-human primate (NHP) models with PD play an essential role in the analysis of PD pathophysiology and behavior symptoms. As impairments of hand dexterity function can affect activities of daily living in patients with PD, research on hand dexterity function in NHP models with chronic PD is essential. Traditional rating scales previously used in the evaluation of animal spontaneous behavior were insufficient due to factors related to subjectivity and passivity. Thus, experimentally designed applications for an appropriate apparatus are necessary. In this study, we aimed to longitudinally assess hand dexterity function using hand dexterity task (HDT) in NHP-PD models. To validate this assessment, we analyzed an alteration in Parkinsonian tremor symptoms, loss of dopaminergic neuron, and positron emission tomography (PET) imaging of dopamine transporters (DAT) in these models. HDT latency significantly increased in NHP-PD models. In addition, a significant inverse correlation between HDT and DAT was identified, but no local bias was found. The correlation with intention tremor symptoms was lower than the resting tremor. In conclusion, the evaluation of HDT may reflect behavioral symptoms of NHP-PD models. Furthermore, HDT was effectively used to experimentally distinguish intention tremors from other tremors.

## Introduction

Symptoms of Parkinson’s disease (PD) caused by loss of dopaminergic neurons are accompanied by movement disorders, including tremors, rigidity, bradykinesia, and akinesia [1, 2]. As impairments of hand dexterity function can affect activities of daily living in patients with PD, such as buttoning a shirt or picking up a fork, recent studies have attempted to identify related pathological processes, such as apraxia, to develop a treatment strategy [3, 4]. Various methods have been developed to assess hand dexterity dysfunction in patients with PD [5–7].

Non-human primate (NHP) of PD models play an essential role in the analysis of PD pathophysiology and behavior symptoms [8]. However, methods that can be applied to NHP models with PD are limited and confined to those based on visual phenotype observations. Traditional rating scales previously used in the evaluation of animal spontaneous behavior were problematic due to factors related to subjectivity and passivity [9]. Observation methods can neither eliminate idiosyncrasies of each observer’s subjective judgment (subjectivity) nor separate bradykinesia from temporary spontaneous laziness of the subjects (passivity). In particular, distinguishing parkinsonian symptoms, such as bradykinesia, from spontaneous laziness in NHP-PD models was challenging [9].

To overcome these challenges, experimentally designed applications of an appropriate apparatus are necessary. An apparatus for the hand dexterity task (HDT) used in experimental NHP models, such as stroke, in recent studies [10–13], can analyze hand dexterity function to measure latency in retrieving food pellets from wells of various sizes and depths. The HDT method has been applied with the expectation that the latency to retrieve food will increase in proportion to the severity of PD symptoms, including bradykinesia, rigidity, and intention tremor in voluntary movements. A performance assessment based on appetite, contrary to passive observation, can evaluate symptoms that occur during voluntary movement and can resolve confounding behavior issues associated with spontaneous laziness.

Furthermore, we believe that an apparatus for the HDT could be used to distinguish intention tremors from other tremors. We suggested that the latency to retrieve food would increase in proportion to the intention tremor severity in voluntary movements. Although Parkinsonian tremors are key symptoms of the disease, understanding the development of tremors remains unclear. In addition, a research on the differentiation between resting and intention tremors in voluntary movements can be a clue for Parkinsonian tremors [14, 15].

In this study, we aimed to longitudinally assess hand dexterity function using the HDT in a previously developed 1-methyl-4-phenyl-1,2,3,6-tetrahydropyridine (MPTP)-induced NHP model for chronic PD [9]. To validate this assessment, we analyzed the alteration of Parkinsonian behavior score, including tremor symptoms and positron emission tomography (PET) imaging of dopamine transporters (DAT), with [^18^F] N-(3-fluoropropyl)-2β-carboxymethoxy-3β-(4-iodophenyl) nortropane (^18^F-FP-CIT).

## Material and Methods

### Experimental animals and MPTP administration

Seven adult female cynomolgus monkeys (Macaca fascicularis) that were developed in a previous study [9] were used in this study. The design for the MPTP (0.2 mg/kg; Sigma-Aldrich, St. Louis, MO, USA) injections was also previously reported [9]. Briefly, MPTP was dissolved in saline. Then four monkeys were intramuscularly injected with MPTP into the left femoral region daily while three monkeys were injected with saline.

### Ethical statement

This animal experiment were approved by the Korea Research Institute of Bioscience and Biotechnology Institutional Animal Care and Use Committee (Approval No. KRIBB-AEC-16068) and conformed to the ARRIVE guidelines [16]. The monkeys originated from the Zhaoqing Laboratory Animal Research Centre (Guangdong Province, China) and were maintained in individual indoor cages at the National Primate Research Center at the Korea Research Institute of Bioscience and Biotechnology (KRIBB), as previously reported [9, 17–21]. In brief, the experimental animals housed in cages faced each other in double columns. So, the animals had visual contact and voice interaction to neighbors without physical contact to avoid the damage to other individuals through the excretion of MPTP metabolites. The dimensions of cages are 60 cm × 80 cm × 80 cm. These cage spaces meet the guidelines of National Institutes of Health in USA. They were fed commercial monkey chow (Teklad 2050™, Envigo, USA) supplemented by various fruits and were given water ad libitum. They were also supplied various rubber and plastic toys and fruits for enrichments. If monkeys do not have an interest in toys, new shape and size of toys were put in cages. Environmental conditions were maintained a temperature of 24 ± 2 degrees Celsius, a relative humidity of 50 ± 5%, and a 12 h light/12 h dark cycle. The attending veterinarian monitored the monkeys’ health in accordance with the recommendations of the Weatherall et al. report on the use of nonhuman primates in research [22]. Animal health monitoring was performed by microbiological tests including B virus, simian retrovirus (SRV), simian immunodeficiency virus (SIV), simian virus 40 (SV40), and simian T-cell lymphotropic virus (STLV) at once a year, as described previously [23].

### Hand dexterity task

A standard HDT instrument was used for the evaluation of hand dexterity dysfunction caused by PD. All monkeys were trained to retrieve 190 mg of pellets (Fruit Crunchies, Bio Serv, USA) from 4 wells of various depths and diameters located in a tray on the left or right side of the apparatus, as explained in detail in previous studies [10–13, 21]. The 4 wells had the following characteristics: Well 1 = narrow and shallow, Well 2 = wide and shallow, Well 3 = wide and deep, Well 4 = narrow and deep; diameter: narrow = 1.9 cm, wide = 2.5 cm; depth: shallow = 0.95 cm, deep = 1.6 cm (Fig 1). The different sizes required different levels of finger dexterity of the monkey to retrieve the food pellets. The testing was conducted before breakfast in order to increase each animal’s appetite. The performance assessment based on appetite made it possible to exclude spontaneous laziness of the animals as a confounding factor. Eight trials were conducted (4 trials per hand) on the trial day once a week, and each trial was limited to 2 min. Three training sessions were conducted for 3 weeks before MTPT or saline injections. The same sequence was maintained for all HDTs. An overall process sequence was recorded using a digital video camera (HDR-CX405, Sony Corp., Japan) fixed on the top of the cage. The captured video was used to measure pellet retrieval latency from each of the four wells. The latency time indicated the time to return from the passage hole with retrieved pellet. Latency cut-off in each well was 15 s. In order to perform a correlation analysis on the HDT and DAT activity at 8, 16 and 24 weeks after MPTP injections, the results of ^18^F-FP-CIT binding potential in PET scan of previous studies were used [9].

**Fig 1.**
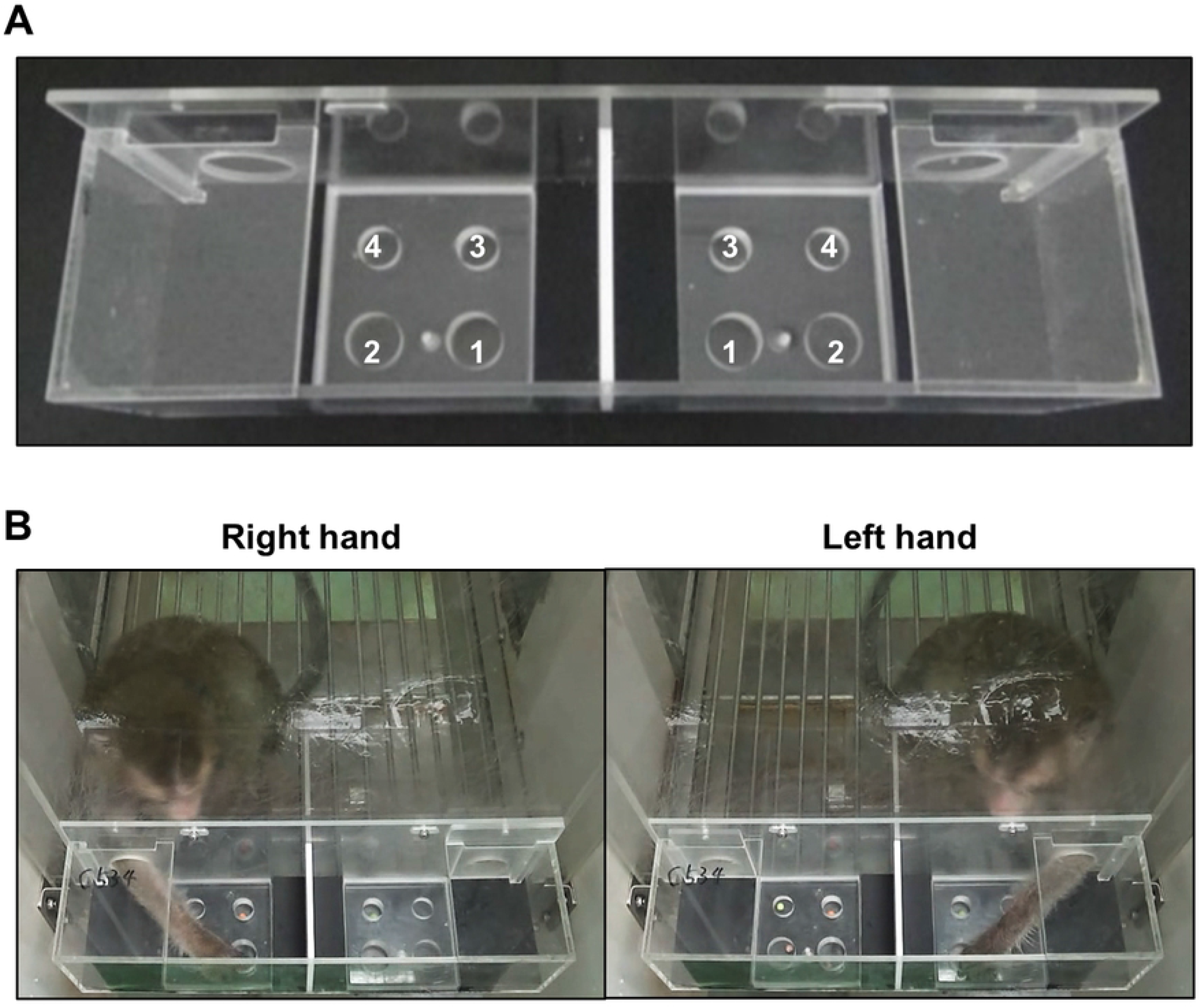
Introduction of the hand dexterity task (HDT) for cynomolgus monkeys. (A) Representative images show an apparatus for the HDT. (B) The captured images during HDT performance for each hand.

### Parkinsonian tremor assessment

The intention tremor assessment was utilized to calculate tremor symptoms during HDT performance using the tremor item *Kurlan scale* [24, 25]. The resting tremor of the monkeys was evaluated according to a rating scale for monkey PD models through a 5 min video recording analyzed every 60 min for a total of 240 min; the tremors were then scored based on the *Kurlan scale*, as in a previous study [9]. The evaluation was performed by three trained examiners who were not informed of the experimental design.

### Statistical analysis

All statistical analyses were performed using the Statistical Package for the Social Sciences for Windows, Version, 18.0 (SPSS Inc., Chicago, IL). Results of the HDT, PET imaging, and tremor scores were analyzed by comparing to baseline, using repeated measures analysis of variance ANOVA. Linear regressions were used to analyze the relationship between HDT, PET imaging results, and tremor scores. *P* < 0.05 was considered statistically significant.

## Results

### Cynomolgus monkeys adapted to HDT for 3 trial days

Three training sessions were conducted for 3 weeks to train the HDT before MTPT or saline injections. All cynomolgus monkeys had significantly adapted to Well 1, 2, and 4 of HDT for 3 trial days. The latency of each wells on three trial days of baseline test recorded was as follows: Well 1 left = 2.01 (± 0.26) sec, Well 1 right = 3.21 (± 0.64) sec, Well 2 left = 0.80 (± 0.08) sec, Well 2 right = 1.23 (± 0.25) sec, Well 3 left = 12.06 (± 1.61) sec, Well 3 right = 12.67 (± 1.52) sec, Well 4 left = 0.95 (± 0.10) sec, Well 4 right = 1.21 (± 0.13) sec (Fig 2). The mean latency of day 3 trial on both hands of each monkey (baseline) was used as a criterion for comparison of the latency to retrieve pellets after MPTP administration on the HDT.

**Fig 2.**
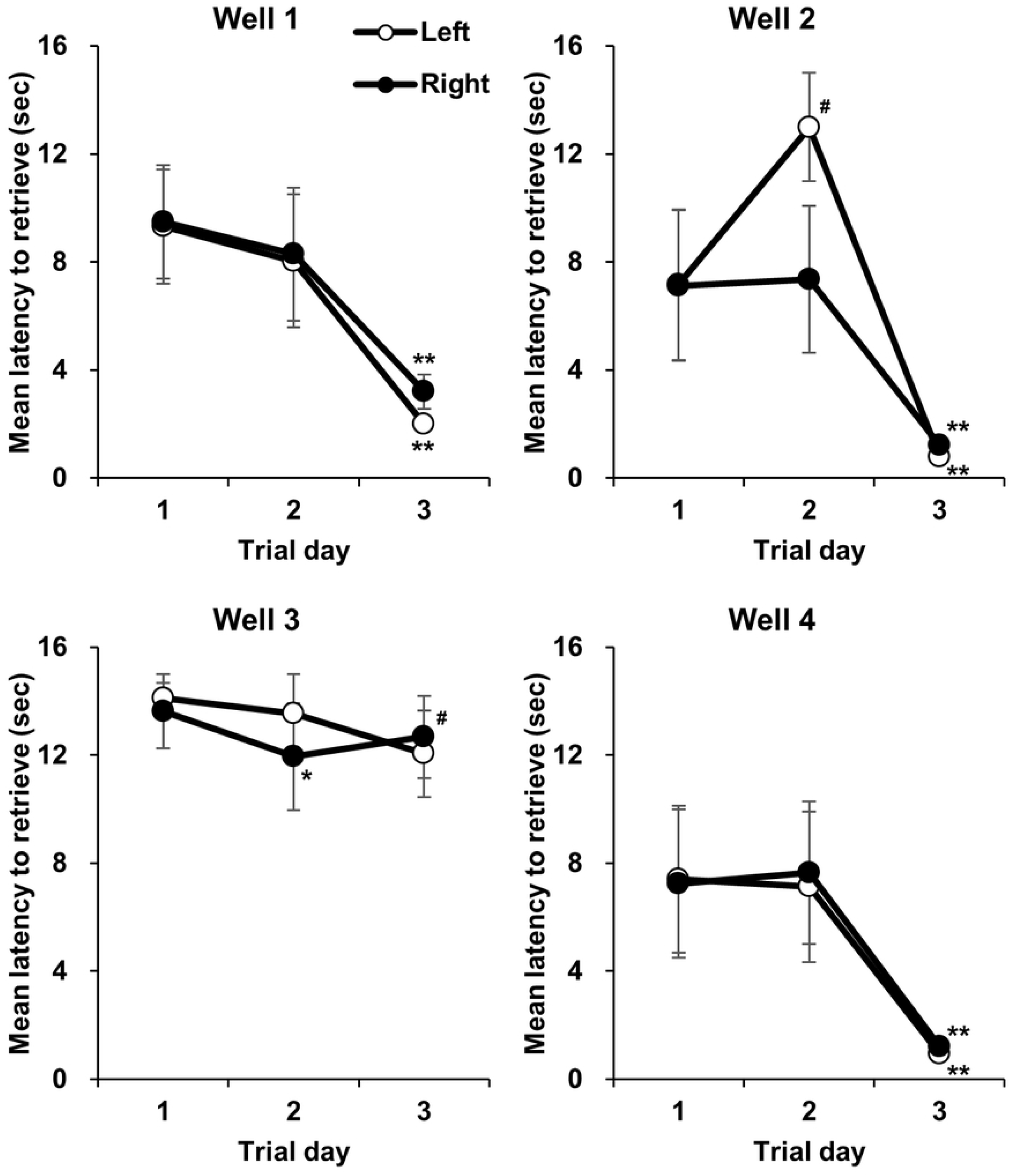
The cynomolgus monkeys adapted to HDT for 3 trial days. All cynomolgus monkeys had adapted to Well 1, 2, and 4 of the HDT for 3 trial days (n = 7). #*P* < 0.05, *\*P* < 0.01 vs. 1 trial day, *\*\*P* < 0.01 vs. 1 and 2 trial day respectively; repeated measures ANOVA.

### Latency identified in the HDT significantly increased after MPTP injections

To confirm functional hand dexterity impairment, we observed the latency of HDT before and after MPTP administration (Fig 3). HDT was used to compare movements from week 1 after the first MPTP administration; latency to retrieve pellets from Well 1 in the HDT task was significantly increased for both hands, compared with baseline, and remained at that level until 24 weeks after the first administration (Fig 3). The tendency of increased latency was also observed in Well 2 and 4; however, latency to retrieve pellets from other wells, except Well 1 of the HDT task, did not significantly change.

**Fig 3.**
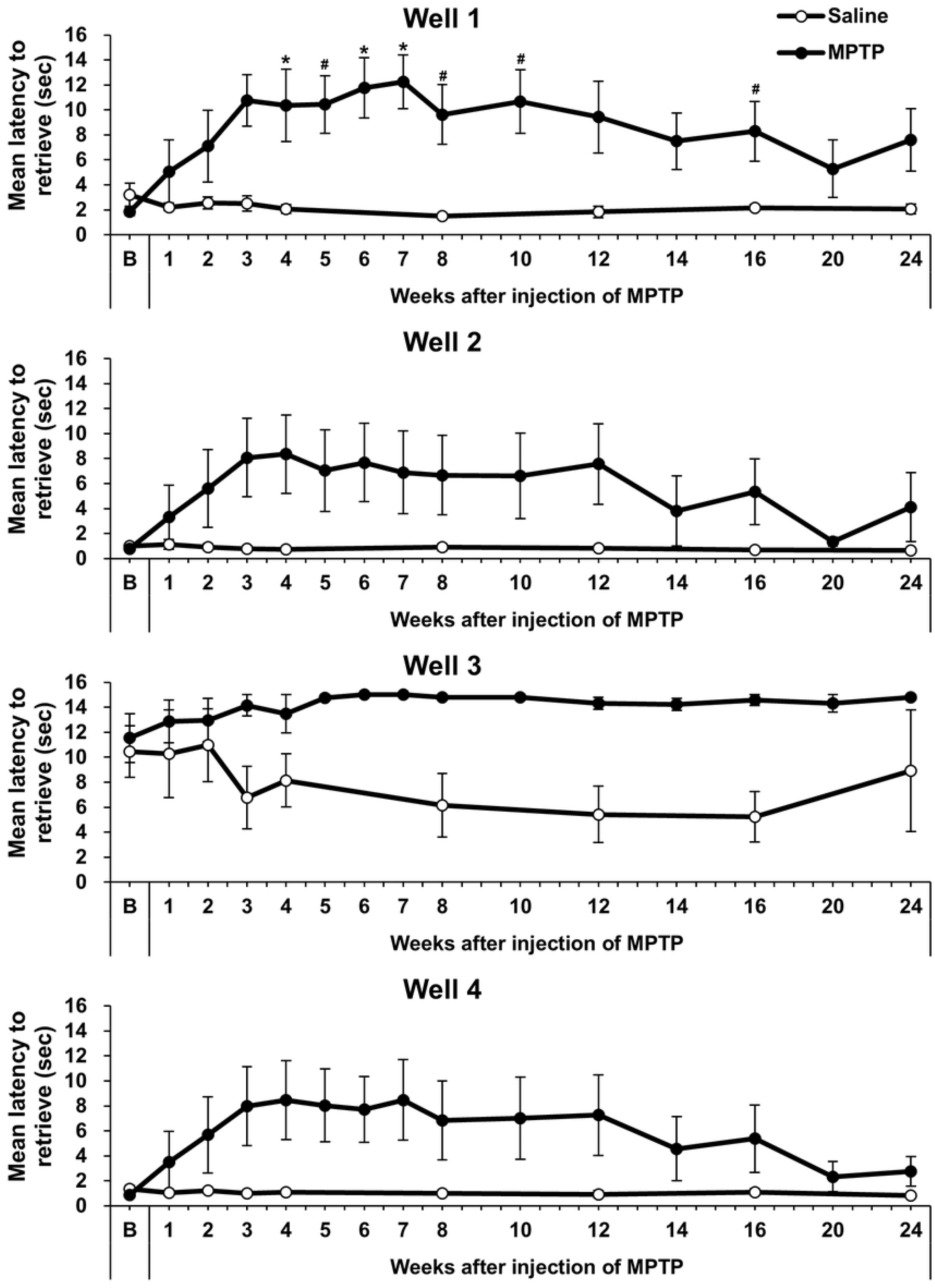
Mean latency to retrieve pellets on the hand dexterity task (HDT). HDT was used to compare movements from week 1 after the first MPTP administration; latency to retrieve pellets from Well 1 in the HDT task was significantly increased for both hands, compared with baseline, and remained at that level until 24 weeks after the first administration. *#P* < 0.05, *\*P* < 0.01 vs. baseline; repeated measures ANOVA. B, baseline.

### Significant inverse correlation between HDT and DAT activity was observed but no local bias was found

To analyze the correlation between HDT and DAT activity, linear regressions were performed using PET results from a previous study [9]. A significant inverse correlation was found between the latency to complete the HDT on Well 1 and the ^18^F-FP-CIT BP in all sub-regions of the striatum on the opposite side (Fig 4, R^2^ = 0.323-0.476, *P* < 0.01); monkeys with fewer striatal DAT tended to have poorer performance, indicating that HDT data reflects the level of striatal DAT. However, no local bias in the correlation between HDT and DAT activity in region of striatum was distinctively observed.

**Fig 4.**
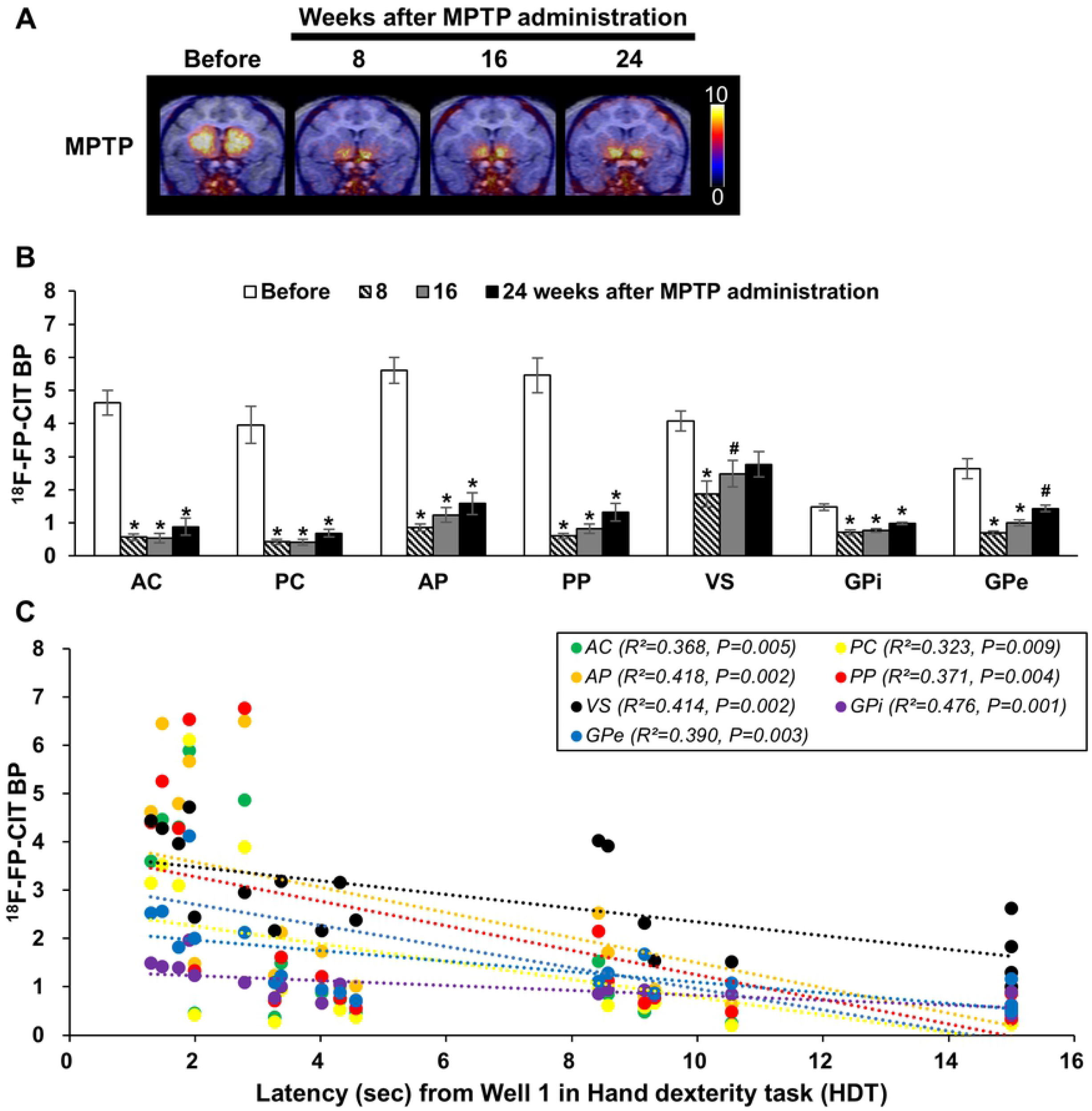
Correlation between hand dexterity task (HDT) and dopamine transporters (DAT) activity. (A) Representative ^18^F-FP-CIT positron emission tomography (PET) images to indicate the DAT at the time points of 8, 16 and 24 weeks after MPTP injections. (B) Histogram representing ^18^F-FP-CIT binding potential (BP) in the MPTP-injection group. (C) Correlations between levels of ^18^F-FP-CIT (BP) with the latency from Well 1 of the hand dexterity task in the MPTP-injection group at all PET time point matched. AC, anterior caudate; PC, posterior caudate; AP, anterior putamen; PP, posterior putamen; VS, ventral striatum; GPi, globus pallidus interna; GPe, globus pallidus external.

### Tremor symptoms during the HDT performance

Neurological examinations of the tremor symptoms were conducted using tremor scores based on the *Kurlan scale* [9, 24, 25]. The tremor assessments were evaluated by analyzing video recordings during the performance of the HDT (S1 and S2 Video) and the spontaneous behavior in individual cages. The tremor score during the spontaneous behavior in individual cages showed continuous tremor symptoms from 3 weeks after the first MPTP administration (Fig 5A). The tremor score recordings during the performance of the HDT showed a tendency to increase, but it did not significantly change. There was a positive correlation between HDT and both the tremor symptom levels (Fig 5, R^2^ = 0.12−0.40, *P* < 0.01). Unexpectedly, the HDT showed higher correlation with the tremor symptom on the spontaneous behavior than the tremor symptom on HDT performance.

**Fig 5.**
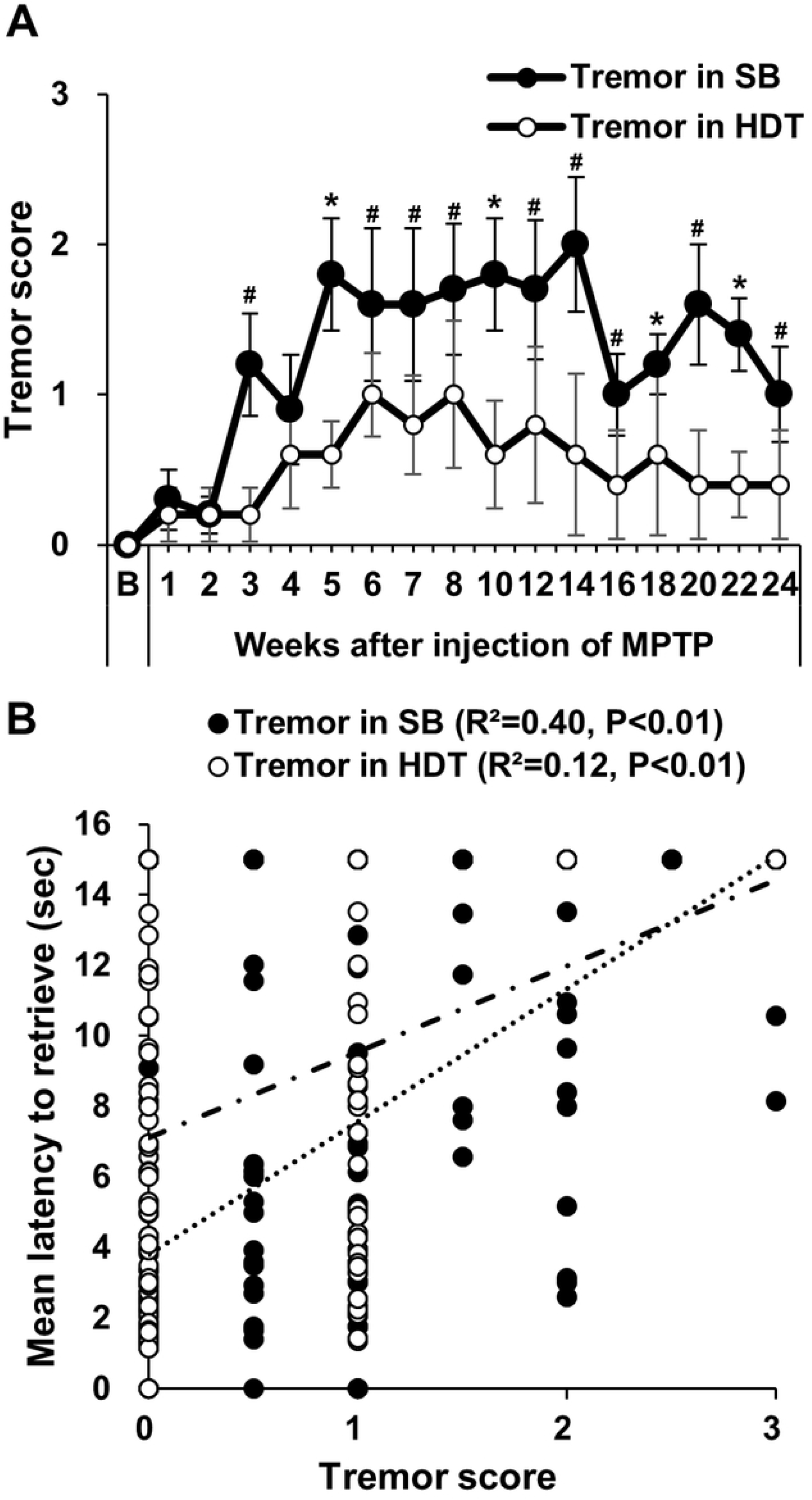
Correlation between hand dexterity task (HDT) and tremor symptoms. (A) The changes of tremors in spontaneous behavior (SB) and hand dexterity task (HDT) after MPTP administration. The tremors scores in SB remained high. (B) Positive correlations between the latency from Well 1 and the tremor score in SB (R^2^ = 0.40, *P*<0.01) and HDT (R^2^ = 0.12, *P*<0.01). *#P* < 0.05, *\*P* < 0.01 vs. baseline; repeated measures ANOVA. SB, Spontaneous behavior; HDT, Hand dexterity task; B, baseline.

## Discussion

The major finding of this study was that Well 1 in the HDT task was sufficiently sensitive to detect impairments of hand dexterity function induced by injection of MPTP. This study suggested that the HDT can investigate the behavior symptoms of NHP-PD models. In addition, HDT can provide verifiable means to investigate parkinsonian hand dexterity function in NHP-PD models.

As individual differences in proficiency and learning ability may affect the outcomes using this apparatus, a comparison with the baseline of the same animal is essential along with the control group of saline-injected animals. A minimum of three training sessions are required for accurate evaluation before the injection of MPTP. After three training sessions, all subjects showed constant latency to retrieve food (Fig 2).

Furthermore, based on voluntary movements, we confirmed that this assessment resolved confounding issues between bradykinesia and spontaneous laziness. All monkeys injected with MPTP performed the HDT despite parkinsonian symptoms, such as bradykinesia and rigidity. The performance assessments based on appetite and foraging habits made it possible to exclude spontaneous laziness in the experimental animals. This method, therefore, is helpful not only to overcome the passivity of traditional observation evaluations but also aid in the investigation of hand dexterity functions in voluntary movements (Fig 3).

In addition, we were able to distinguish intention tremor from other tremors during HDT performance (S1 and S2 video). Although controversial, movements that occur during appetite-based HDT tasks can be considered goal-directed movements. Hence, we considered that tremor symptoms during the HDT tasks indicate intention tremor. Several clinical studies in patients with PD explained that resting and postural tremors during involuntary movements are more prominent than intention tremor during goal-directed movements [26–29]. However, studies pertaining to tremor symptoms mechanism remain insufficient, due to difficulty in animal model analysis [30, 31]. Although several quantitative studies of tremor symptom in NHPs PD model were performed [32–35], it is challenging to distinguish rest tremor and other tremors for these animal models experimentally [31]. We believe that an attempt to distinguish type of tremor symptoms using HDT task for NHP models provides an important conceptual advance in the study of NHP-PD models and tremor symptoms.

In recent study of the marmoset MPTP model, the evidence of fine motor skills impairments and postural head tremor symptom were provided after MPTP injection using the object retrieval task with barrier detour (ORTBD) [36]. The ORTBD is similar to the HDT, but with a difference. Both tasks are the reward based behavioral testing system, but the HDT is more intuitive compared to ORTBD bypassing the barrier. The cognitive and problem solving skills are required to solve the detour part in the ORTBD, as evidenced in the studies [36–38]. Whereas, the HDT is able to evaluate the impairments of hand dexterity function independently.

We anticipated that it would be helpful to analyze impaired hand dexterity function in correlating analysis between the regional dopaminergic neuronal activity in the basal ganglia and tremor symptom of PD. No local bias in the correlation between HDT and DAT activity in the region of striatum was distinctively observed, although this study showed that HDT latency weakly indicates levels of the DAT activity induced by MPTP (Fig 4). Furthermore, with this method it was also expected that the latency to retrieve food would increase in proportion to the severity of intention tremor in voluntary movements. However, correlation with intention tremor symptoms was lower than the resting tremor in MPTP induced NHP-PD models (Fig 5). The reason for this phenomenon is that the MPTP-induced PD models with impaired dopamine systems may indicate low relevance to intention tremor symptoms [30, 39]. According to previous studies, Parkinsonian tremor symptoms controlled by altered cerebral activity in the globus pallidus and the cerebello-thalamo-cortical circuits, distinct from the nigrostriatal pathway, could result in impaired hand dexterity function [40, 41]. In addition, a recent study described how a combination of both the basal ganglia and cerebellum plays a crucial role in tremor phenotypes [31]. The serotonergic system and gamma-aminobutyric acid (GABA)-ergic systems have also been implicated in PD [42–44]. Further precise comparisons of the behavior symptoms and neuropathology between NHP models may enhance better understanding of the mechanism of rest tremor in PD.

## Conclusions

In conclusion, HDT can be helpful in the investigation of behavior symptoms of NHP-PD models. In addition, HDT enables experimental distinction between intention tremors and other tremors, and the correlation of HDT with intention tremor symptoms was lower than the resting tremor in NHP-PD model induced by MPTP.

## Acknowledgments

None

## Supporting information

**S1 Video. The recorded video during HDT performance for each hand before MPTP injections.**

**S2 Video. The recorded video during HDT performance for each hand at 8 weeks after MPTP injections.**

## References

1. Kalia LV, Lang AE. Parkinson’s disease. The Lancet. 2015;386(9996):896–912. doi: 10.1016/s0140-6736(14)61393-3.

2. Poewe W, Seppi K, Tanner CM, Halliday GM, Brundin P, Volkmann J, et al. Parkinson disease. Nat Rev Dis Primers. 2017;3:17013. doi: 10.1038/nrdp.2017.13. PubMed PMID: 28332488.

3. Vanbellingen T, Kersten B, Bellion M, Temperli P, Baronti F, Muri R, et al. Impaired finger dexterity in Parkinson’s disease is associated with praxis function. Brain Cogn. 2011;77(1):48–52. doi: 10.1016/j.bandc.2011.06.003. PubMed PMID: 21775040.

4. Foki T, Pirker W, Klinger N, Geissler A, Rath J, Steinkellner T, et al. FMRI correlates of apraxia in Parkinson’s disease patients OFF medication. Exp Neurol. 2010;225(2):416–22. doi: 10.1016/j.expneurol.2010.07.019. PubMed PMID: 20659452.

5. Wenzelburger R, Raethjen J, Loffler K, Stolze H, Illert M, Deuschl G. Kinetic tremor in a reach-to-grasp movement in Parkinson’s disease. Mov Disord. 2000;15(6):1084–94. PubMed PMID: 11104190.

6. Dahdal P, Meyer A, Chaturvedi M, Nowak K, Roesch AD, Fuhr P, et al. Fine Motor Function Skills in Patients with Parkinson Disease with and without Mild Cognitive Impairment. Dement Geriatr Cogn Disord. 2016;42(3-4):127–34. doi: 10.1159/000448751. PubMed PMID: 27643700.

7. Pradhan SD, Brewer BR, Carvell GE, Sparto PJ, Delitto A, Matsuoka Y. Assessment of fine motor control in individuals with Parkinson’s disease using force tracking with a secondary cognitive task. J Neurol Phys Ther. 2010;34(1):32–40. doi: 10.1097/NPT.0b013e3181d055a6. PubMed PMID: 20212366.

8. Emborg ME. Nonhuman primate models of Parkinson’s disease. ILAR J. 2007;48(4):339–55. doi: 10.1093/ilar.48.4.339. PubMed PMID: 17712221.

9. Seo J, Lee Y, Kim BS, Park J, Yang S, Yoon HJ, et al. A non-human primate model for stable chronic Parkinson’s disease induced by MPTP administration based on individual behavioral quantification. J Neurosci Methods. 2019;311:277–87. doi: 10.1016/j.jneumeth.2018.10.037. PubMed PMID: 30391524.

10. Moore TL, Killiany RJ, Pessina MA, Moss MB, Rosene DL. Assessment of motor function of the hand in aged rhesus monkeys. Somatosens Mot Res. 2010;27(3):121–30. doi: 10.3109/08990220.2010.485963. PubMed PMID: 20653499; PubMed Central PMCID: PMCPMC6504938.

11. Moore TL, Killiany RJ, Pessina MA, Moss MB, Finklestein SP, Rosene DL. Recovery from ischemia in the middle-aged brain: a nonhuman primate model. Neurobiol Aging. 2012;33(3):619 e9-e24. doi: 10.1016/j.neurobiolaging.2011.02.005. PubMed PMID: 21458887; PubMed Central PMCID: PMCPMC3145025.

12. Moore TL, Pessina MA, Finklestein SP, Kramer BC, Killiany RJ, Rosene DL. Recovery of fine motor performance after ischemic damage to motor cortex is facilitated by cell therapy in the rhesus monkey. Somatosens Mot Res. 2013;30(4):185–96. doi: 10.3109/08990220.2013.790806. PubMed PMID: 23758412; PubMed Central PMCID: PMCPMC6503838.

13. Moore TL, Pessina MA, Finklestein SP, Killiany RJ, Bowley B, Benowitz L, et al. Inosine enhances recovery of grasp following cortical injury to the primary motor cortex of the rhesus monkey. Restor Neurol Neurosci. 2016;34(5):827–48. doi: 10.3233/RNN-160661. PubMed PMID: 27497459; PubMed Central PMCID: PMCPMC6503840.

14. Helmich RC, Hallett M, Deuschl G, Toni I, Bloem BR. Cerebral causes and consequences of parkinsonian resting tremor: a tale of two circuits? Brain. 2012;135(Pt 11):3206–26. doi: 10.1093/brain/aws023. PubMed PMID: 22382359; PubMed Central PMCID: PMCPMC3501966.

15. Loane C, Wu K, Bain P, Brooks DJ, Piccini P, Politis M. Serotonergic loss in motor circuitries correlates with severity of action-postural tremor in PD. Neurology. 2013;80(20):1850–5. doi: 10.1212/WNL.0b013e318292a31d. PubMed PMID: 23596065; PubMed Central PMCID: PMCPMC3908354.

16. Kilkenny C, Browne WJ, Cuthill IC, Emerson M, Altman DG. Improving bioscience research reporting: the ARRIVE guidelines for reporting animal research. PLoS Biol. 2010;8(6):e1000412. doi: 10.1371/journal.pbio.1000412. PubMed PMID: 20613859; PubMed Central PMCID: PMCPMC2893951.

17. Yeo HG, Lee Y, Jeon CY, Jeong KJ, Jin YB, Kang P, et al. Characterization of Cerebral Damage in a Monkey Model of Alzheimer’s Disease Induced by Intracerebroventricular Injection of Streptozotocin. J Alzheimers Dis. 2015;46(4):989–1005. doi: 10.3233/JAD-143222. PubMed PMID: 25881906.

18. Lee Y, Kim YH, Park SJ, Huh JW, Kim SH, Kim SU, et al. Insulin/IGF signaling-related gene expression in the brain of a sporadic Alzheimer’s disease monkey model induced by intracerebroventricular injection of streptozotocin. J Alzheimers Dis. 2014;38(2):251–67. doi: 10.3233/JAD-130776. PubMed PMID: 23948941.

19. Park J, Seo J, Won J, Yeo HG, Ahn YJ, Kim K, et al. Abnormal Mitochondria in a Non-human Primate Model of MPTP-induced Parkinson’s Disease: Drp1 and CDK5/p25 Signaling. Exp Neurobiol. 2019;28(3):414–24. doi: 10.5607/en.2019.28.3.414. PubMed PMID: 31308800; PubMed Central PMCID: PMCPMC6614070.

20. Seo J, Park J, Kim K, Won J, Yeo HG, Jin YB, et al. Chronic Infiltration of T Lymphocytes into the Brain in a Non-human Primate Model of Parkinson’s Disease. Neuroscience. 2020;431:73–85. doi: 10.1016/j.neuroscience.2020.01.043. PubMed PMID: 32036014.

21. Kim K, Jeon H-A, Seo J, Park J, Won J, Yeo H-G, et al. Evaluation of cognitive function in adult rhesus monkeys using the finger maze test. Applied Animal Behaviour Science. 2020;224:104945. doi: 10.1016/j.applanim.2020.104945.

22. Weatherall D. The use of non-human primates in research. 2006.

23. Jeong HS, Lee SR, Kim JE, Lyoo IK, Yoon S, Namgung E, et al. Brain structural changes in cynomolgus monkeys administered with 1-methyl-4-phenyl-1,2,3,6-tetrahydropyridine: A longitudinal voxel-based morphometry and diffusion tensor imaging study. PLoS One. 2018;13(1):e0189804. doi: 10.1371/journal.pone.0189804. PubMed PMID: 29320500; PubMed Central PMCID: PMCPMC5761839.

24. Imbert C, Bezard E, Guitraud S, Boraud T, Gross CE. Comparison of eight clinical rating scales used for the assessment of MPTP-induced parkinsonism in the Macaque monkey. J Neurosci Methods. 2000;96(1):71–6. PubMed PMID: 10704673.

25. Kurlan R, Kim MH, Gash DM. Oral levodopa dose-response study in MPTP-induced hemiparkinsonian monkeys: assessment with a new rating scale for monkey parkinsonism. Mov Disord. 1991;6(2):111–8. doi: 10.1002/mds.870060205. PubMed PMID: 2057003.

26. Puschmann A, Wszolek ZK. Diagnosis and treatment of common forms of tremor. Semin Neurol. 2011;31(1):65–77. doi: 10.1055/s-0031-1271312. PubMed PMID: 21321834; PubMed Central PMCID: PMCPMC3907068.

27. Bhidayasiri R. Differential diagnosis of common tremor syndromes. Postgrad Med J. 2005;81(962):756–62. doi: 10.1136/pgmj.2005.032979. PubMed PMID: 16344298; PubMed Central PMCID: PMCPMC1743400.

28. Deuschl G, Bain P, Brin M. Consensus statement of the Movement Disorder Society on Tremor. Ad Hoc Scientific Committee. Mov Disord. 1998;13 Suppl 3:2–23. doi: 10.1002/mds.870131303. PubMed PMID: 9827589.

29. Dirkx MF, Zach H, Bloem BR, Hallett M, Helmich RC. The nature of postural tremor in Parkinson disease. Neurology. 2018;90(13):e1095–e103. doi: 10.1212/WNL.0000000000005215. PubMed PMID: 29476038; PubMed Central PMCID: PMCPMC5880634.

30. Emborg ME, Tetrud JW, Moirano J, McLaughlin WW, Bankiewicz KS. Rest tremor in rhesus monkeys with MPTP-induced parkinsonism. Front Biosci. 2003;8:a148–54. doi: 10.2741/1088. PubMed PMID: 12700090.

31. Pan MK, Ni CL, Wu YC, Li YS, Kuo SH. Animal Models of Tremor: Relevance to Human Tremor Disorders. Tremor Other Hyperkinet Mov (N Y). 2018;8:587. doi: 10.7916/D89S37MV. PubMed PMID: 30402338; PubMed Central PMCID: PMCPMC6214818.

32. Bergman H, Raz A, Feingold A, Nini A, Nelken I, Hansel D, et al. Physiology of MPTP tremor. Mov Disord. 1998;13 Suppl 3:29–34. doi: 10.1002/mds.870131305. PubMed PMID: 9827591.

33. Oiwa Y, Eberling JL, Nagy D, Pivirotto P, Emborg ME, Bankiewicz KS. Overlesioned hemiparkinsonian non human primate model: correlation between clinical, neurochemical and histochemical changes. Front Biosci. 2003;8:a155–66. doi: 10.2741/1104. PubMed PMID: 12957824.

34. German DC, Dubach M, Askari S, Speciale SG, Bowden DM. 1-Methyl-4-phenyl-1,2,3,6-tetrahydropyridine-induced parkinsonian syndrome in Macaca fascicularis: which midbrain dopaminergic neurons are lost? Neuroscience. 1988;24(1):161–74. doi: 10.1016/0306-4522(88)90320-x. PubMed PMID: 3259295.

35. Deutch AY, Elsworth JD, Goldstein M, Fuxe K, Redmond DE, Jr., Sladek JR, Jr., et al. Preferential vulnerability of A8 dopamine neurons in the primate to the neurotoxin 1-methyl-4-phenyl-1,2,3,6-tetrahydropyridine. Neurosci Lett. 1986;68(1):51–6. doi: 10.1016/0304-3940(86)90228-4. PubMed PMID: 3487756.

36. Choudhury GR, Daadi MM. Charting the onset of Parkinson-like motor and non-motor symptoms in nonhuman primate model of Parkinson’s disease. PLoS One. 2018;13(8):e0202770. doi: 10.1371/journal.pone.0202770. PubMed PMID: 30138454; PubMed Central PMCID: PMCPMC6107255 alter our adherence to PLOS ONE policies on sharing data and materials.

37. Taylor JR, Elsworth JD, Roth RH, Sladek JR, Jr., Redmond DE, Jr. Cognitive and motor deficits in the acquisition of an object retrieval/detour task in MPTP-treated monkeys. Brain. 1990;113 (Pt 3):617–37. doi: 10.1093/brain/113.3.617. PubMed PMID: 2364263.

38. McEntire CR, Choudhury GR, Torres A, Steinberg GK, Redmond DE, Jr., Daadi MM. Impaired Arm Function and Finger Dexterity in a Nonhuman Primate Model of Stroke: Motor and Cognitive Assessments. Stroke. 2016;47(4):1109–16. doi: 10.1161/STROKEAHA.115.012506. PubMed PMID: 26956259.

39. Tetrud JW, Langston JW. Tremor in MPTP-induced parkinsonism. Neurology. 1992;42(2):407–10. doi: 10.1212/wnl.42.2.407. PubMed PMID: 1736174.

40. Dirkx MF, den Ouden HE, Aarts E, Timmer MH, Bloem BR, Toni I, et al. Dopamine controls Parkinson’s tremor by inhibiting the cerebellar thalamus. Brain. 2017. doi: 10.1093/brain/aww331. PubMed PMID: 28073788.

41. Caligiore D, Helmich RC, Hallett M, Moustafa AA, Timmermann L, Toni I, et al. Parkinson’s disease as a system-level disorder. npj Parkinson’s Disease. 2016;2:16025. doi: 10.1038/npjparkd.2016.25.

42. Lang EJ, Sugihara I, Llinas R. GABAergic modulation of complex spike activity by the cerebellar nucleoolivary pathway in rat. J Neurophysiol. 1996;76(1):255–75. doi: 10.1152/jn.1996.76.1.255. PubMed PMID: 8836223.

43. Galvan A, Wichmann T. GABAergic circuits in the basal ganglia and movement disorders. Prog Brain Res. 2007;160:287–312. doi: 10.1016/S0079-6123(06)60017-4. PubMed PMID: 17499121.

44. Pagano G, Politis M. Molecular Imaging of the Serotonergic System in Parkinson’s Disease. Int Rev Neurobiol. 2018;141:173–210. doi: 10.1016/bs.irn.2018.08.002. PubMed PMID: 30314596.

